# Response of Regulatory Genetic Variation in Gene Expression to Environmental Change in *Drosophila melanogaster*

**DOI:** 10.1101/2020.06.23.163501

**Authors:** Wen Huang, Mary Anna Carbone, Richard F. Lyman, Robert H. H. Anholt, Trudy F. C. Mackay

## Abstract

The genetics of phenotypic responses to changing environments remains elusive. Using whole genome quantitative gene expression as a model, we studied how the genetic architecture of regulatory variation in gene expression changed in a population of fully sequenced inbred *Drosophila melanogaster* strains when flies developed at different environments (25 °C and 18 °C). We found a substantial fraction of the transcriptome exhibited genotype by environment interaction, implicating environmentally plastic genetic architecture of gene expression. Genetic variance in expression increased at 18 °C relative to 25 °C for most genes that had a change in genetic variance. Although the majority of expression quantitative trait loci (eQTLs) for the gene expression traits in the two environments were shared and had similar effects, analysis of the environment-specific eQTLs revealed enrichment of binding sites for two transcription factors. Finally, although genotype by environment interaction in gene expression could potentially disrupt genetic networks, the co-expression networks were highly conserved across environments. Genes with higher network connectivity were under stronger stabilizing selection, suggesting that stabilizing selection on expression plays an important role in promoting network robustness.

## Introduction

Organisms living in fluctuating environments or entering novel environments must possess mechanisms to cope with environmental changes. One such mechanism is to change expressed phenotypes in response to different environments, a phenomenon called phenotypic plasticity (Price et al., 2003). Alternatively, organisms may develop homeostatic mechanisms to cushion the effect of environmental fluctuations without changing their phenotypes. The maintenance of homeostasis is important for organismal fitness as it protects organisms from detrimental effects. The definition of environment can be broad, ranging from cell types, tissues, and physiological states, to diseases, external stimuli, and climate; all of which are known to cause plastic changes or lack thereof in certain phenotypes.

In addition to environmental factors, phenotypes can also respond to genetic perturbations in a plastic or homeostatic manner, which characterizes the potential of an organism to express phenotypes when genes mutate. In a population of genetically diverse individuals, the extent of genetic variation of a phenotype measures the overall sensitivity of individuals to mutations segregating in the population.

Importantly, the state of plasticity or homeostasis, with respect to either genetic or environmental variation, is not necessarily static and can be modified by both genetic and environmental factors. A classic example is the heat shock protein system, particularly *Hsp90*, whose expression is environmentally plastic and increases under thermal stress, but buffers phenotypic changes induced by mutations to maintain homeostasis (Rutherford and Lindquist, 1998), a process termed canalization (Waddington, 1942). The opposite of canalization – decanalization – describes the change from a homeostatic state to a plastic one, which allows phenotypic expression of genetic and/or environmental variation (Geiler-Samerotte et al., 2019; Gibson and Wagner, 2000). The dynamics of genetic variation (variance across different genotypes) and environmental variation (variance across different environments) may be controlled by different mechanisms. For example, although the histone variant H2A.Z is a capacitator for environmental variation (Levy and Siegal, 2008), its presence in the yeast genome does not increase robustness to mutations (Richardson et al., 2013).

Change in genetic variation across environments is one of the many forms of genotype by environment interaction (GxE). GxE can be interpreted equivalently either as variable genetic architecture across environments or as variable environmental plasticity across genotypes, depending on what factor is chosen as the context. GxE has important implications in quantitative trait variation and evolution. It is important for maintenance of genetic variation (Gillespie and Turelli, 1989). It is pervasive in plants and animals and influences domestication (Doust et al., 2014) and genetic improvement (Des Marais et al., 2013; Rauw and Gomez-Raya, 2015). GxE is also of paramount importance to realize personalized medicine such as individualized drug therapy (Eichelbaum et al., 2006).

Gene expression is a unique class of quantitative traits that are under genetic control and that exhibit both plasticity and homeostasis (López-Maury et al., 2008). Because of its sheer number and annotations of biological functions, gene expression can serve as an important model for quantitative traits. In this study, we sought to understand the response of the genetic architecture of gene expression by exposing the sequenced inbred lines of the *Drosophila melanogaster* Genetic Reference Panel to a low temperature treatment. Temperature is one of the most wide ranging environmental factors an organism can experience and must manage, and previous studies have shown that there is genetic variation in plasticity of fitness in response to low temperature in *Drosophila melanogaster* (Fry et al., 1998). We measured whole-genome gene expression levels and combined them with a previous data set that quantified gene expression of the same lines reared at standard ambient temperature (Huang et al., 2015). This experimental design whereby the same standing genetic variation is subjected to different thermal environments enabled us to address three fundamental questions. First, how does genetic variance of gene expression change when the environment changes? Second, is environmental plasticity of gene expression heritable; or equivalently, does heritable regulatory variation change in response to environmental change? If so, what are the locations and effects of the quantitative trait loci (QTLs) that exhibited such variability and response to environments? And finally, how does the response of regulatory genetic variation in gene expression change the architecture of co-expression networks?

## Results

### Identification of transcriptional units by RNA sequencing and estimation of gene expression by tiling microarrays

We used a two-stage procedure (Figure 1a) to measure whole-genome gene expression profiles for adult flies independently raised at 18 °C and 25 °C for 185 *Drosophila melanogaster* Genetic Reference Panel (DGRP) lines (Mackay et al., 2012). In the first stage, we defined regions in the genome that expressed detectable RNA levels by sequencing pooled polyadenylated RNA from all DGRP lines for each of the two sexes at the two temperatures separately (Table S1, Figure S1). After alignment of RNA sequence reads to the reference transcriptome and genome, followed by transcript model reconstruction, we merged known and newly discovered transcript models from all conditions to obtain 21,873 gene models with non-overlapping constitutive exons (Figure 1a).

**Figure 1.**
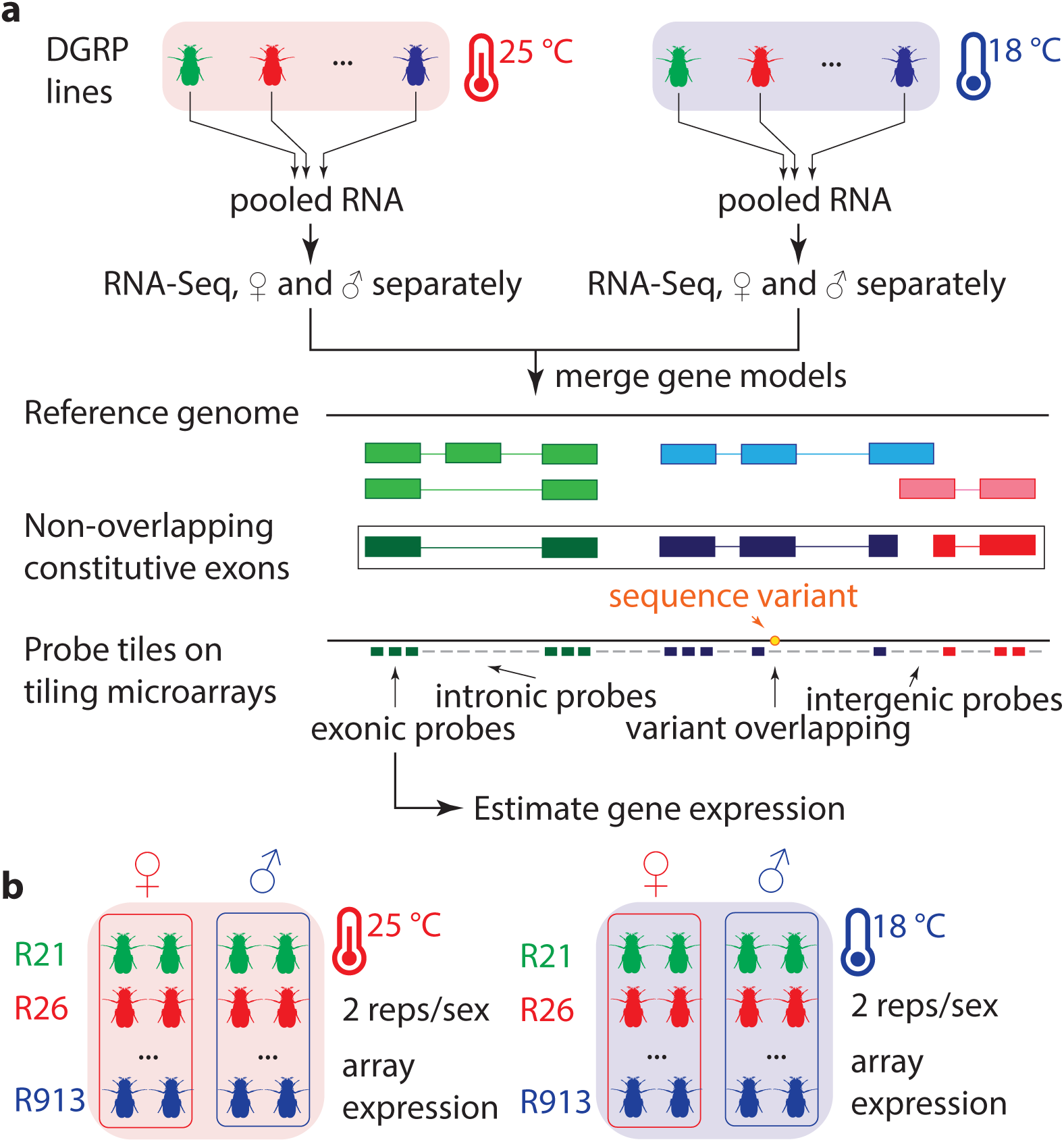
Experimental design and gene expression estimation strategy. (**a**) RNA extracted from adult flies raised at 25 °C and 18 °C from all DGRP lines were pooled and sequenced to define a set of non-overlapping constitutive exons. Genome tiling arrays were then used to estimate gene expression, summarizing expression based on intensities of probes that fall entirely within constitutive exons and do not overlap with common sequence variants. (**b**) DGRP lines were raised at 25 and 18 °C and RNA was extracted from adult flies for two biological replicates per sex within each temperature. Gene expression was estimated using genome tiling arrays.

In the second stage, we used genome tiling arrays to estimate gene expression for the DGRP lines with two replicates per line, sex, and temperature (Figure 1b). After removing probes that overlapped with common non-reference alleles, we were able to estimate expression for 20,691 genes, among which about 24% (*n* = 4,943) were unannotated novel transcribed regions (NTRs). Our subsequent global analyses did not differentiate between annotated genes and NTRs, except when performing gene set enrichment analyses when annotations were needed. We filtered out six samples (none involved both replicates of the same condition) that were outliers based on scaled expression within each sex and temperature (Figure S2) and removed undesired batch effects using surrogate variable analysis (Leek and Storey, 2007) on normal quantile transformed expression within each sex. The adjustment appeared to be effective because known batches such as array scan date was effectively captured by the derived surrogate variables (Figure S3). Subsequent analyses began with these adjusted expression values.

### Canalization and decanalization of genetic variance in gene expression

Because we collected the data at 18 °C and 25 °C separately, the effects of temperature and batch were confounded. Therefore we cannot make inferences on the effect of temperature on the mean change of gene expression, which has been extensively investigated in other studies (Chen et al., 2015; Levine et al., 2011). To characterize the patterns of genetic variance in gene expression in the two thermal environments, we first partitioned the variance in gene expression into the between-line genetic component 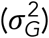 and the remaining within-line micro-environmental component 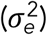. In females, there were 4,912 genes with significant genetic variance at 18 °C, and 3,002 at 25 °C at an FDR = 0.05 (Table S2), among which 2,505 were shared between both environments. In males, there were 5,315 and 4,278 genes with significant genetic variance at 18 °C and 25 °C respectively, including 3,339 in common (Table S2).

The marked difference in the numbers of genes with significant genetic variance at 18 °C and 25 °C in both sexes was intriguing, and could be due to either an overall decrease in environmental variance or an increase in genetic variance at 18 °C. We therefore tested for variance heterogeneity for the genetic and environmental components. In both sexes, 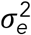 was relatively stable across the two temperatures for the majority of genes (Figure 2a-d, Table S2). In contrast, the difference in 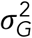 was far more pronounced (Figure 2a-d, Table S2). This pattern of variance heterogeneity was almost identical when we did not adjust for infection status with the symbiont *Wolbachia* bacteria that affects approximately half of the lines (Figure S4), consistent with the previous observation that *Wolbachia* infection does not substantially impact gene expression (Everett et al., 2020; Huang et al., 2015).

**Figure 2.**
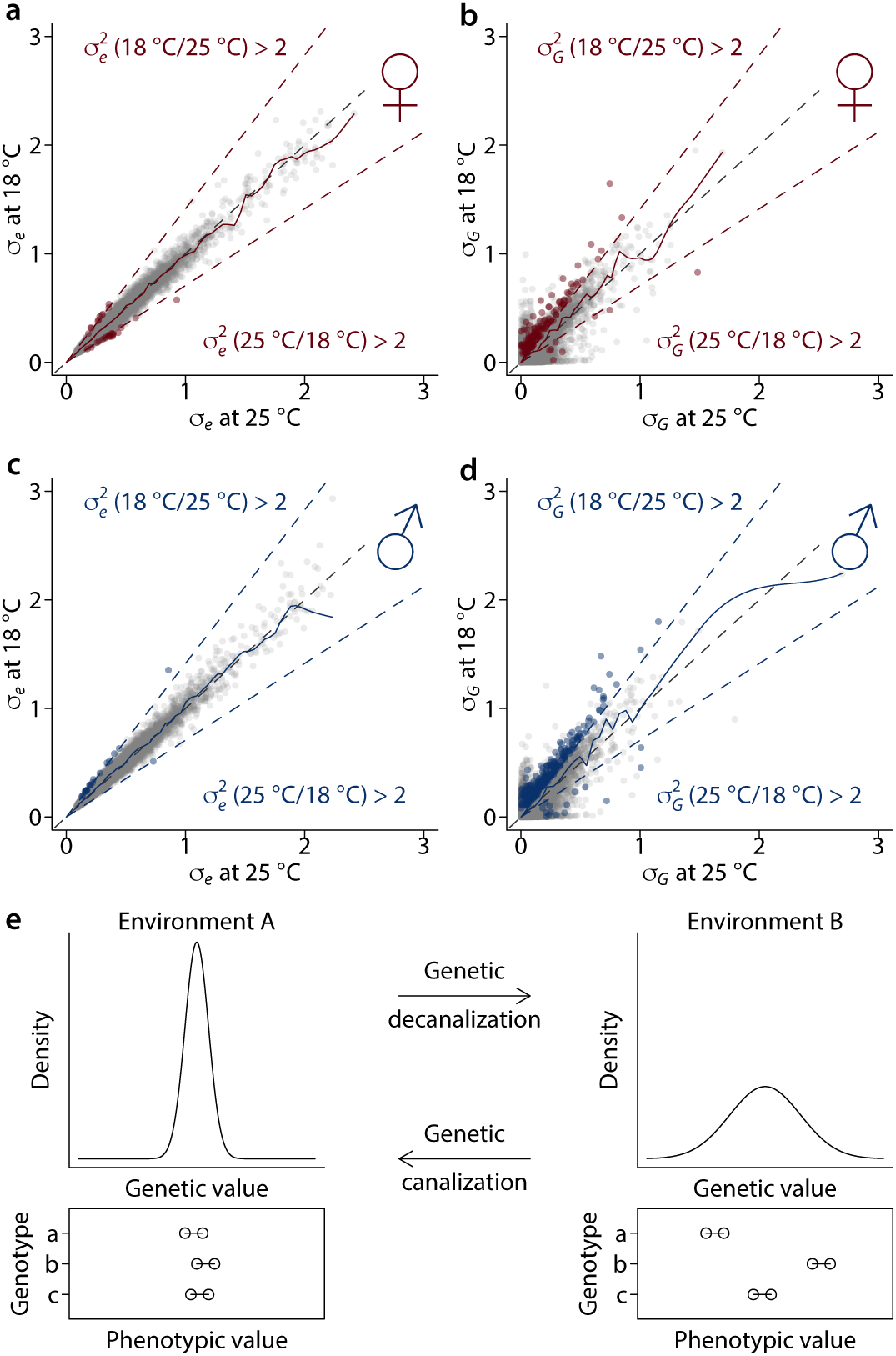
Decanalization and canalization of genetic variance of gene expression at 18 °C. (**a**) Comparison of environmental variation of gene expression between 18 and 25 °C in females. The two outer dashed lines indicate two-fold changes, while the diagonal line indicates no change. Note that the axes were standard deviations while the two-fold change was assessed for variances. A LOESS smooth line (dark solid line) is added with a span of 0.001 at the scale of the plotted axis. Red points indicate genes with signiicant variance heterogeneity for 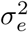 (FDR = 0.05) by at least two fold changes. (**b**) Comparison of genetic variation of gene expression between 18 and 25 °C in females. The dashed lines indicate two-fold changes, while the diagonal line indicates no change. A LOESS smooth line (dark solid line) is added with a span of 0.001 at the scale of the plotted axis. Red points indicate genes with significant variance heterogeneity for 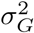 (FDR = 0.05) by at least two fold and signiicant 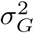 (FDR = 0.05) in the environment with higher genetic variance. (**c**) and (**d**), comparison of environmental and genetic variances in males, where blue points indicate genes with signiicant variance heterogeneity. (**e**) Plots illustrating the dynamics of genetic variances (genetic decanalization/canalization) between two environments. The density plots show the distributions of genetic values for individuals in a population in two environments (A and B). Below the density plots are line graphs showing phenotypic values where points connected by lines are biological replicates from the same genotypes to illustrate micro-environmental variance.

We define genetic decanalization as increased genetic variance in gene expression, and genetic canalization as decreased genetic variance in an environment relative to an arbitrary baseline (Figure 2e). Our tests for variance heterogeneity at the two temperatures revealed both genetic decanalization and canalization for gene expression relative to 25 °C when flies developed at 18 °C, depending on specific genes considered (Figure 2). Interestingly, genetic decanalization at 18 °C relative to 25 °C was more prevalent than canalization. There were 149 genes in females that had significantly different genetic variance at 18 °C than 25 °C by at least two fold, among which 141 were genetically decanalized and only 8 were canalized (Table S2). The same was true in males, where 264 genes were decanalized and 15 were canalized at 18 °C. We found little evidence for preferential genetic canalization or decanalization of the expression of genes involved in particular functions. Using gene set enrichment analysis (GSEA), only three broad Gene Ontology (GO) terms were found to be significantly (FDR = 0.05) enriched for genes whose genetic variance was decanalized or canalized at 18 °C, including the structural constituent of chitin-based cuticle that was enriched for decanalized genes in females and odorant binding and DNA-binding transcription factor activity enriched for canalized genes in males (Table S3, Figure S5).

### Response of regulatory genetic variation in gene expression to environmental change

The change in genetic variance upon exposure to environmental treatment is a special form of genotype by environment interaction (GxE). To understand the genetic basis of genotype by environment interaction in gene expression, we asked whether there was variation in gene expression that could be attributed to GxE. There are several equivalent ways to describe this phenomenon, each with a different perspective. The first is a largely statistical description, which can be graphically illustrated by reaction norms. In the reaction norm representation, the presence of GxE causes otherwise parallel lines depicting environmental plasticity to cross (Figure 3a). GxE may or may not be accompanied by an environmental effect when averaged across individuals. Importantly, with the same phenotypic scale across environments, a change in genetic variance will cause a statistical presentation of GxE (Figure 3b) though the reverse is not necessarily true. Alternatively, GxE is equivalent to environmentally responsive differences between genotypes (Figure 3c). If environmental plasticity is genetically variable, its variation between DGRP lines characterizes how heritable the plasticity is (Figure 3a). Finally, if an environment is able to modify the allelic effects of QTLs controlling the phenotype (Barreiro et al., 2011; Fairfax et al., 2014; Lee et al., 2014), significant GxE indicates that the genetic architecture of the phenotype is environmentally responsive.

**Figure 3.**
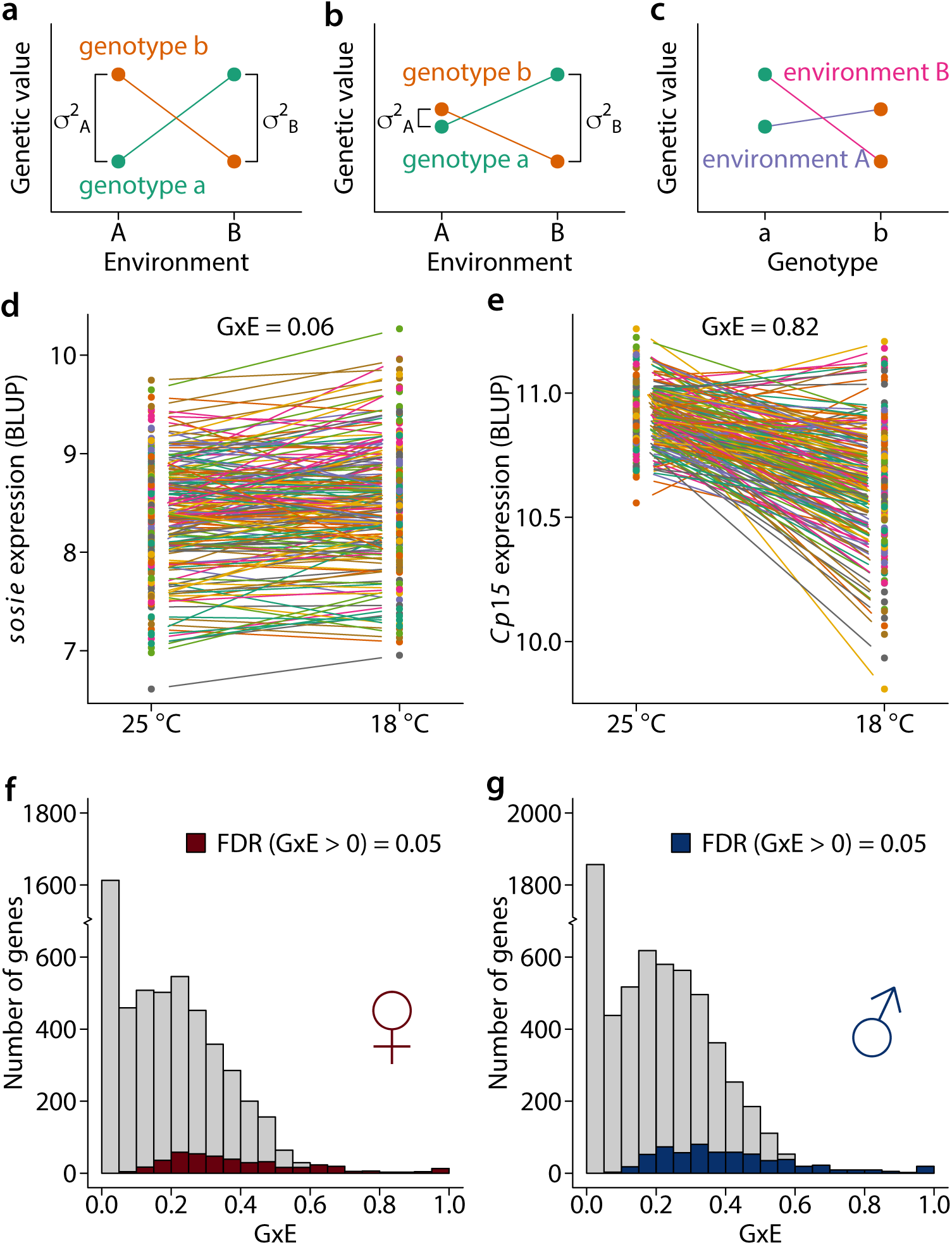
Genotype by environment interaction (GxE) of gene expression. (**a**) Reaction norms between environment A and B for two genotypes a and b. The lines connect points of the same colors represent the different genotypes. In this example, there is no change in genetic variance. (**b**) No mean effect of temperature but significant GxE and change in genetic variance. (**c**) The same data as in **a** but plotted with respect to difference between genotypes across two environments. (**d**) Example of a gene (*sosie*) in females with low degree of GxE. (**e**) Example of a gene (*Cp75*) with a high degree of GxE in females. GxE is deined as 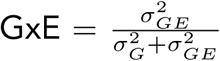. (**f**) Distribution of GxE for genes in females, where signiicant genes were marked by dark red color. (**g**) Distribution of GxE for genes in males, where signiicant genes were marked by dark blue color.

To identify genes that showed significant genotype by environment interaction, we pooled the data from both environments and partitioned the variance in gene expression into components due to a common genetic effect shared by both temperatures 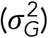 and due to GxE 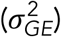. Remarkably, among the 5,248 genes in females and 6,327 genes in males that had at least some significant genetic component (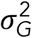 or 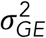), 424 (8%) and 619 (10%) respectively had significant GxE at an FDR = 0.05 (Table S2, Figure 3d,e). The genetic architecture of regulatory variation of these genes was thus variable or environmentally responsive between the two thermal environments. Equivalently, the environmental plasticity of these genes in response to low temperature was therefore heritable. Of these genes, 66 and 110 were also significant for variance heterogeneity between the two environments in females and males respectively. GSEA revealed little evidence of GxE or the lack of it concentrating in particular biological functions as only a very limited number of broad GO terms were enriched for higher or lower degree of GxE, including tricarboxylic acid cycle and translation initiation enriched for larger GxE in males and protein serine/threonine kinase activity enriched for smaller GxE in males at FDR = 0.05 (Table S4).

### Mapping response of regulatory genetic variation to environmental change

We have previously shown that we can map expression quantitative trait loci (eQTLs) for a substantial fraction of genes that are genetically variable in the same population (Huang et al., 2015). To understand the environmental response of the genetic architecture of gene expression, we mapped eQTLs at both 18 °C and 25 °C among 1,891,697 common (minor allele frequency > 0.05) variants and compared the locations and effects of the eQTLs. In females, there were 793 and 511 genes with at least one mapped eQTL (FDR = 0.05), constituting approximately 16% and 17% of the genetically variable genes at 18 °C and 25 °C respectively. In males, we mapped eQTLs (FDR = 0.05) for 1,086 (20%) and 808 (19%) genes at 18 °C and 25 °C, respectively. To refine the eQTL-gene association models and account for linkage disequilibrium between DNA variants, we performed forward model selection such that variants meeting the FDR thresholds were added in the order of their association with gene expression conditional on existing variants in the model until no variant could be added at *P* < 1 x 10^−5^. This procedure resulted in between 1 and 5 eQTLs for each gene, with the majority (74%) of genes containing only one eQTL (Table S5).

Through the mapped eQTLs and their estimated effects and locations, we made four comparisons in order to understand the response of the regulatory genetic variation to environmental change in the Drosophila transcriptome. First, we asked if there were shared or environment-specific eQTLs, regardless of the signs and magnitudes of effects. In females, there were 2,407 and 497 genes that had environment-specific genetic variance at 18 °C and 25 °C respectively, among which 222 (9%) and 41 (8%) contained mapped eQTLs (Figure 4a). In males, 1,976 and 939 genes were genetically variable only at 18 °C and 25 °C, respectively; 250 (13%) and 116 (12%) of which contained mapped eQTLs (Figure 4b). These environment-specific eQTLs represent regulatory genetic variation that responds (‘inactive’ versus ‘active’) to the specific environmental change. In addition, of the 2,505 genetically variable genes at both 18 °C and 25 °C in females, 571 and 470 genes had mapped eQTLs respectively (Figure 4a). Among the 329 genes with at least one eQTL in both environments (sharing of the dark purple boxes in Figure 4a), 303 (92%) had at least one shared eQTL. A similar result was obtained for males, where 459 (93%) out of 494 genes with eQTLs in both environments shared common eQTLs (Figure 4b). These results indicate that when genes contained eQTLs in both environments, the majority of them shared eQTLs.

**Figure 4.**
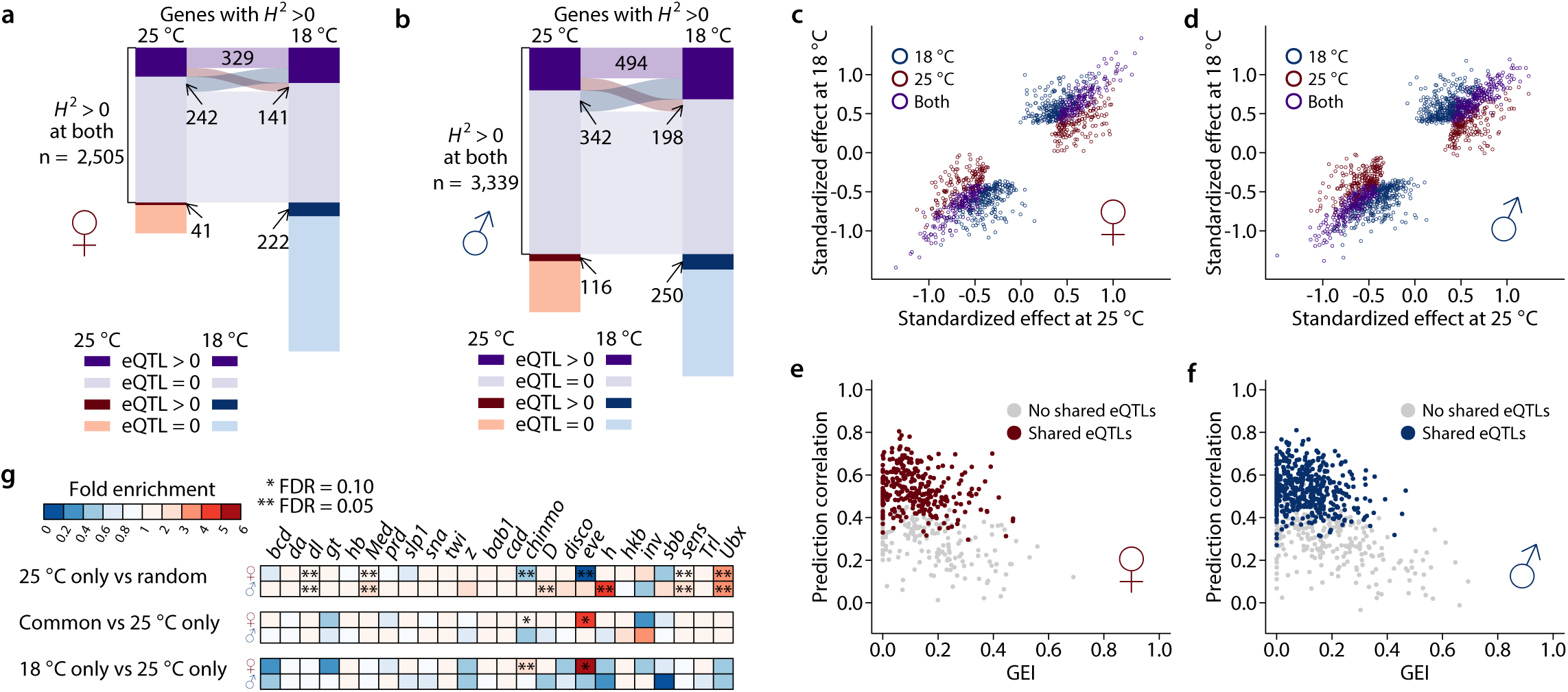
Plasticity of regulatory variation in gene expression. (**a** – **b**) Sharing of genes with eQTLs between 25 °C and 18 °C in females (**a**) and males (**b**). The numbers indicate genes only. Each colored box represents the number of genes in each category, with the category indicated on the bottom left, where eQTL > 0 indicates genes with mapped eQTLs while eQTL = 0 indicates genes without mapped eQTLs. Boxes with the same colors (dark and light purple) are genes that are genetically variable at both temperatures. The size of sharing is proportional to the width of the ribbons connecting the boxes. (**c** – **d**) eQTL effect comparison between 25 and 18 °C in females (**c**) and males (**d**). (**e** – **f**) Cross-environment prediction of gene expression using eQTLs in females (**e**) and males (**f**). Gene expression at 18 °C was predicted based on eQTLs mapped at 25 °C and the prediction was evaluated by Pearson correlation between the observed and predicted expression. The prediction correlation was plotted against GEI. (**g**) Enrichment/depletion of transcription factor binding sites overlap among eQTLs. The enrichment was computed based on ratio over the mean of overlap among 10,000 randomly drawn sets of eQTLs with the same numbers across the three classes (25 °C only, 18 °C only, and common). *P*-value was computed based on the number of iterations that was more extreme than the observed overlap and adjusted using the Benjamini-Hochberg approach.

Second, we compared the estimated single variant effects of eQTLs at 18 °C and 25 °C among the genes genetically variable at both temperatures. Of the 1,181 eQTL-gene pairs in females, 294 (25%) were specific to 25 °C, 436 (37%) were specific to 18 °C, and 451 (38%) were common to both (Figure 4c). Of the 1,740 eQTL-gene pairs in males, 412 (24%) were specific to 25 °C, 630 (36%) were specific to 18 °C, and 698 (40%) were common to both (Figure 4d). However, these classifications depended on significance thresholds. To enable a more quantitative comparison, we estimated single variant genetic effects regardless of their statistical significance as long as it was significant in at least one environment. As expected, for eQTLs that were shared by both environments, their effects were large and highly similar (Figure 4c,d). On the other hand, eQTLs that were specific to either environment had much larger effects in the environment where they were detected (Figure 4c,d). Those eQTLs whose effects were different between environments also contributed to the environmental response of the regulatory variation. Interestingly, almost all eQTLs had effects of the same sign in both environments, suggesting that GxE or the environmental response of regulatory genetic variation for the majority of genes was likely a result of change in the magnitudes of effects rather than signs.

Third, we asked if using the estimated eQTL effects in one environment could predict gene expression in the other. Conserved regulatory genetic variation would lead to better prediction accuracy. We predicted gene expression at 18 °C for each gene using mapped eQTLs and their effects at 25 °C. As expected, the prediction accuracy correlated well with GxE. Genes with higher GxE, who also tended to not share eQTLs in the two environments, were poorly predicted across environments (Figure 4e,f).

Finally, to probe the nature of the regulatory genetic effects on plasticity, we compared the degree of overlap between the eQTLs of different classes (shared or environment-specific) with known transcription factor binding sites in the Drosophila genome. In both females and males, eQTLs tended to overlap with transcription factor binding sites in general (Figure 4g, 25 °C only vs random), consistent with their localization to the proximity of transcription start and end sites (Huang et al., 2015). Two developmentally important transcription factors, *chinmo* and *eve*, were significantly (FDR = 0.05) depleted among eQTLs specific to 25 °C in females (Figure 4g). This led to the over-representation of eQTLs common to both environments (FDR = 0.10) or specific to 18 °C (FDR = 0.05 for *chinmo* and 0.10 for *eve*) overlapping the binding sites of these two transcription factors (Figure 4g). Therefore, *chinmo* and *eve* may be involved in the environment specific regulation of gene expression in females.

### Robust co-expression networks in the presence of environmentally responsive regulatory genetic variation

It has been well documented that genes form co-expression networks in which genes executing similar functions have correlated expression levels, either across different tissues of the same individuals, or across individuals in a population (Ardlie et al., 2015). Importantly, perturbations to co-expression networks can lead to expressed phenotypes such as diseases (de la Fuente, 2010; Lea et al., 2019). In a population sample, if there was no genetic variation in the plasticity of gene expression, the co-expression networks constructed based on correlations between genes would remain the same even if plasticity was widespread but constant across individuals. In contrast, GEI or plasticity of regulatory variation may lead to a change in the network structure.

To examine the robustness (or lack thereof) of the co-expression networks in the presence of heritable transcriptome plasticity, we identified co-expression modules using weighted correlation network analysis (WGCNA) in both temperatures and compared the module memberships (Figure 5a,b). We observed strong preservation of network structures. For example, in females, the largest module at 25 °C included 386 genes enriched for basic biological processes such as neurogenesis, mitotic spindle organization, and translation, among others (Table S6). Of these genes, 90% (*n* = 346) were also in the same module at 18 °C (Figure 5a). Using a permutation based approach (Ritchie et al., 2016), several measures of network preservation were found to be highly significant (Table S7).

**Figure 5.**
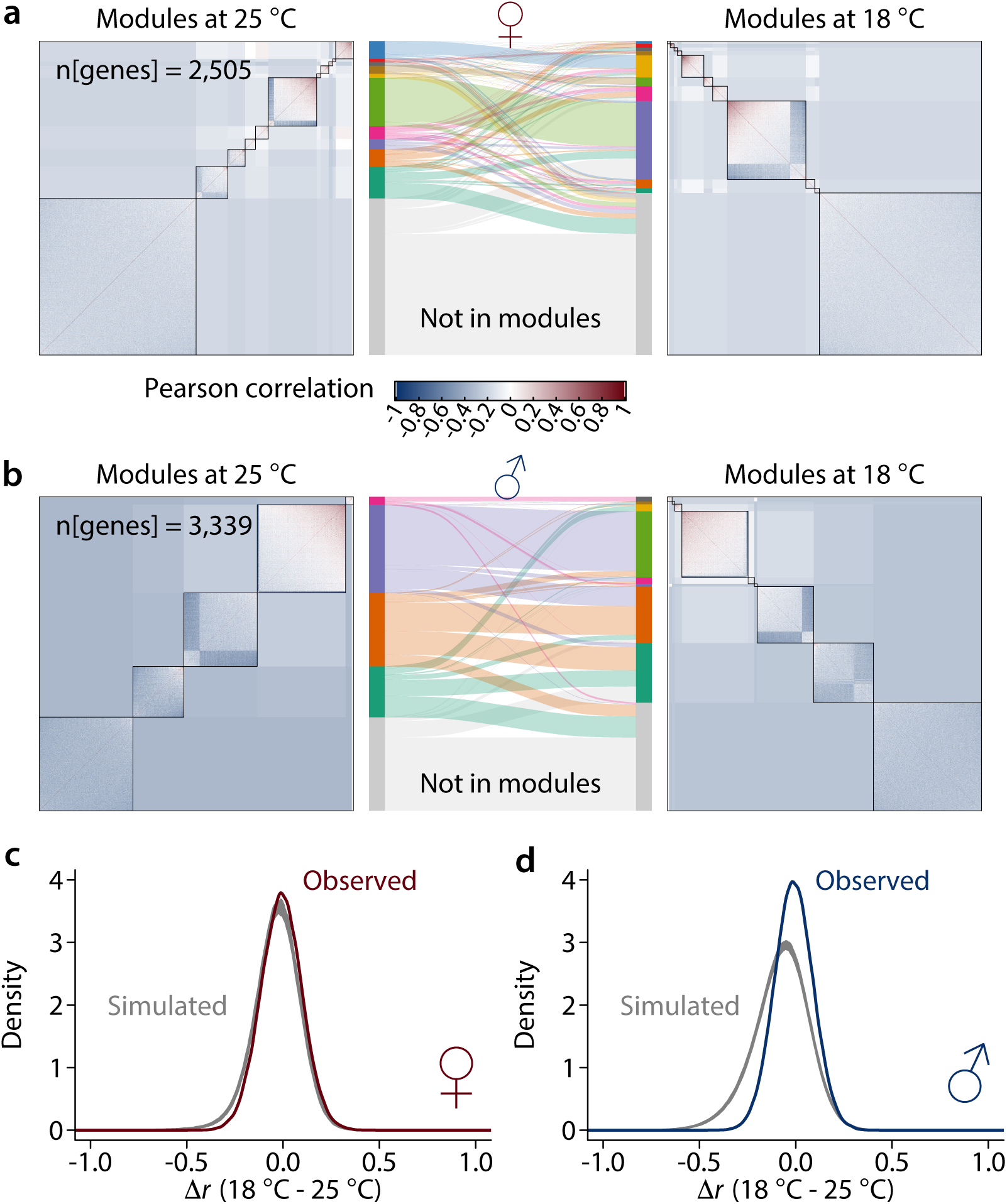
Robust co-expression networks in the presence of plastic regulatory variation. (**a** – **b**) Module preservation between 25 °C and 18 °C in females (**a**) and males (**b**). (**c** – **d**) Density plots for Δ*r* between 18 and 25 °C where the grey lines are based on 1,000 simulations while the dark colored line is the observed in females (**c**) and males (**d**). The test for signiicance was based on variance of the distribution of Δ*r*. The null hypothesis was equal variance while the alternative hypothesis was smaller variance.

However, given that only approximately 10% of genes showed GxE or plasticity of regulatory variation, a high degree of preservation was expected, especially for genes that are required to maintain basic organismal functions. To test the trivial explanation that the network structure preservation is expected given the observed extent of GxE in the transcriptome, we employed a simulation-based approach to derive the expected distribution of changes in correlation under the null hypothesis that the plastic changes in the regulatory variation of genes were independent from each other (Figure 5c,d). We found evidence that the changes in the correlation among genes were smaller than if all genes were responding to the environment change independently. Between the genes with significant GxE, the change in their correlation with each other was significantly smaller than expected (*P* < 0.001 based on 1,000 simulations), more so in males than in females (Figure 5c,d). This result indicates that mechanisms exist to ensure that plasticity of regulatory variation in gene expression remains coordinated between genes in the face of environmental perturbations, thus preserving the co-expression networks.

If the envrionmental response of regulatory genetic variation is not independent but coordinated, genes that are highly correlated with a large number of genes (‘hub’ genes) may be of particular importance in preserving the co-expression networks. We postulated that expression of these genes may be under stronger constraints. We tested this hypothesis by relating the connectivity of genes with the strength of stabilizing selection estimated in a previous study (Huang et al., 2016). We defined the connectivity of a gene as the mean of the absolute correlation of the gene with all other genes, and the strength of stabilizing selection as the ratio of mutational variance (*V*_*m*_) over standing genetic variation (*V*_*g*_). Remarkably, there was a highly significant positive correlation (Spearman *r* = 0.29 and *P* = 1.79e-39 in females and *r* = 0.27 and *P* = 1.23e-44 in males) between gene’s network connectivity and the strength of stabilizing selection in both sexes (Figure 6a,b), indicating that the robustness of co-expression networks is under stabilizing selection. The correlation was slightly stronger for genes without GxE (Figure 6a,b). Though conceivable and suspected in many cases, few studies have experimentally observed this result.

**Figure 6.**
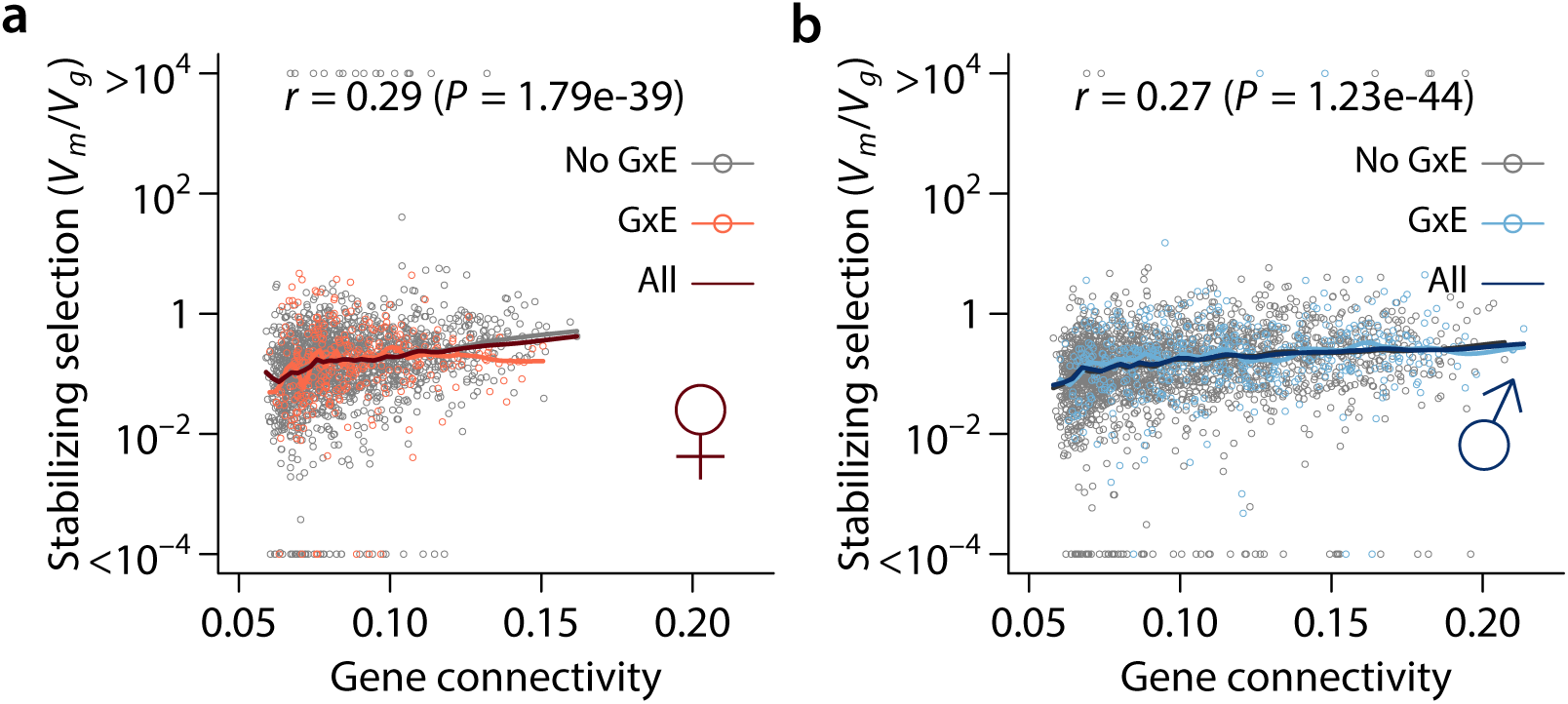
Positive correlation between gene connectivity and strength of stabilizing selection. Strength of stabilizing selection as measured by the ratio between mutational variance (*V m*) and standing genetic variance (*V g*) is plotted against gene connectivity (mean |*r*|) in females (**a**) and (**b**). The dark linees are LOESS smooth lines with a span of 0.1. The colors of the points and lines indicate genes with no GxE, GxE, or all genes. Spearman’s correlation (*r*) and the corresponding *P*-values were also indicated for all genes. In females (**a**), the Spearman’s correlation (*P*) value) was 0.30 (*P* = 1.06*e* − 37) for genes without GxE, 0.19 (*P* = 5.11*e* − 4) for genes with GxE. In males (**a**), the Spearman’s correlation (*P*) value) was 0.27 (*P* = 5.06*e* − 35) for genes without GxE, 0.21 (*P* = 3.89*e* − 6) for genes with GxE.

## Discussion

Using a simple but powerful design, we provided a comprehensive characterization of the response of the regulatory genetic variation of the Drosophila transcriptome to environmental change. Specifically, an inbred line reference population enabled us to use the same genotypes for the alternative treatments and obtain biological replicates. This was instrumental in partitioning the variance in gene expression and providing a global characterization of the extent of genetic canalization or decanalization and GxE. Although we found many more genes genetically decanalized at 18 °C relative to the baseline at 25 °C, this depended on the choice of the baseline environment as well as the environmental history of this population that had shaped its genetic variation. Nevertheless, the genetic variance of hundreds of genes changed between the two thermal environments, clearly indicating that even mild environmental fluctuations can be a potent agent in exposing or masking genetic variation in quantitative trait phenotypes. In addition, approximately 8-10% of genetically variable genes exhibited GxE in just two environments. These genes may be subject to differential selection when they experience heterogeneous environments, contributing to the maintenance of genetic variation (Gillespie and Turelli, 1989). Because only two environments were considered in this study, the extent of GxE may be much more widespread than can be detected and all inferences drawn from this study should be interpreted in the specific context of the two temperatures.

We compared the list of genes exhibiting GxE in the present study with genes that showed evidence of adaptive divergence or GxE between tropical and temperate geographical locations. Among the 619 genes significant for GxE in males in this study, 166 (Table S8) were previously reported to be differentially expressed between flies of tropical and temperate origins (Hutter et al., 2008; Zhao et al., 2015) or had genotype by environment interaction (Levine et al., 2011). However, more formal integration across studies with proper experimental design and statistical inference is needed to understand the role of canalization and decanalization and GxE in general in adaptive evolution, which remains controversial (Partridge and Barton, 2000).

We mapped eQTLs at 18 °C and 25 °C and compared their locations and effects. While there were many environment-specific or plasticity eQTLs, when eQTLs can be mapped in both environments the majority of eQTLs were in fact shared between the two environments with similar effects (Figure 4a-d). Cross-environment prediction of gene expression using mapped eQTLs was poor in the presence of GxE, a caveat that must be considered when predicting gene expression based on mapped eQTLs in reference populations (Gamazon et al., 2015). We found two transcription factors (*chinmo* and *eve*) whose binding sites were enriched in eQTLs in 18 °C relative to 25 °C specific eQTLs, suggesting that they may be involved in regulating the plasticity of gene expression when flies were exposed to 18 °C. The specific mechanisms by which these *cis* regulatory elements change how flies respond to different temperatures require further investigation. Differential transcription factor binding by the developmentally dynamic *chinmo* and *eve* may modify the development of sensory neurons such that flies sense environmental temperature differently (Alpert et al., 2020).

We observed strong preservation of the co-expression network structure between the two environments. A trivial explanation for this observation could be that since the majority of genes did not show significant GxE, the correlation among these genes was preserved. We developed a simulation-based test to specifically test whether the robustness of co-expression networks was expected under the null hypothesis that the plastic responses to low temperature was independent for each of the significant genes. This hypothesis was rejected, indicating that the plastic responses by the genes were coordinated to a certain extent. This would be consistent with a model where the responses of many genes were secondary to a smaller number of primary first responders. Our identification of two transcription factors in females whose binding sites were enriched in eQTLs in 18 °C (common between 25 °C and 18 °C or specific to 18 °C) was also consistent with this model. A robust co-expression network involving many genes would also promote polygenicity of organismal traits or fitness if we consider the expression of genes as mediators of mutational effects on organismal traits. This would be consistent with the ominigenic model of complex quantitative traits (Boyle et al., 2017). Indeed, adaptation to novel thermal environments has been found to be highly polygenic (Barghi et al., 2019). However, further experiments are needed to define a working model.

Importantly, we found genes with higher network connectivity with other genes were also under stronger stabilizing selection (Figure 6). Furthermore, larger and more conserved modules were enriched for genes involved in basic biological processes (Figure 5a,b, Table S6). Previous theoretical work has suggested that stabilizing selection was important in promoting the evolution of network robustness (Espinosa-Soto, 2016). Our result, one of the few (Wagner, 2000), provides empirical support for the role of stabilizing selection in evolving network robustness.

## Methods

### Sample preparation

For the low temperature treatment, all DGRP strains were reared on cornmeal-molasses-agar medium at 18 °C with 60-75% relative humidity and a 12-h light/dark cycle. For each DGRP line, we collected 25 or 40 mated 3-5 day old flies to constitute a biological replicate for females and males respectively. Collection was performed between 1 and 3PM consistently to account for circadian rhythm in gene expression and the flies were immediately frozen in liquid nitrogen before they were sorted. We collected two biological replicates per sex for each of the DGRP lines (*n* = 185 for this study).

### RNA-Sequencing and analysis

We sequenced pooled polyadenylated RNA of flies reared at 18 °C exactly as previously described (Huang et al., 2015). RNA-Seq data for flies reared at 25 °C were downloaded from GEO (GSE67505) and data from the present study were deposited to SRA (PRJNA615927). Although all analyses were identical, we re-analyzed the 25 °C data together with the new 18 °C data for consistency and improvement in sensitivity by merging transcript reconstructions across environments. Briefly, RNA-Seq reads were aligned to the reference transcriptome (FlyBase release 5.57) and the genome (BDGP5) using tophat2 (Kim et al., 2013) (2.0.13) for each sex and temperature separately. Summary of alignments including statistics of input, successful alignments, and reads overlapping various genomic features are provided in Table S1. Alignments were assembled using cufflinks (Trapnell et al., 2013) (2.2.1) into transcript models, which were merged across multiple conditions. To enable subsequent expression profiling using microarrays, we removed segments of exons that were either specific to only a subset of splice isoforms or overlapped from either strand by multiple genes to arrive at a set of non-overlapping constitutive exons for each gene (Figure 1).

### Microarray data acquisition and processing

We acquired and processed microarray data as previously described (Huang et al., 2015). The 25 °C data were downloaded from ArrayExpress (E-MTAB-3216) and analyzed together with the new 18 °C data (E-MTAB-8953). We removed probes that mapped to multiple genomic locations, overlapped with variants whose non-reference allele frequency in the 185 DGRP lines exceeded 0.05, or did not entirely fall within the constitutive exons as defined above. Probe hybridization intensity was corrected for background hybridization (Wu et al., 2004) and quantile normalized (Bolstad et al., 2003) within each sex but across temperature, which was motivated by strong effects of sex on gene expression (Huang et al., 2015) but relatively minor temperature effects as observed in the RNA-Seq data (Figure S1), and an attempt to partially account for batch effects. Probe expression was summarized into gene expression by median polish, which is robust to rare outliers. Finally, within each sex, the expression for each gene was normal quantile transformed with mean equal to the median and standard deviation equal to the median absolute deviation multiplied by a factor of 1.4824. It is important to note that because the 18 and 25 °C data were collected at different times, batch is completely confounded with temperature, and therefore we could not make any inference about the overall effect of temperature on gene expression. Nonetheless, because the arrays were randomized within each temperature, we could still make inferences on GEI after the temperature effect and other latent batch effects were accounted for. We removed unwanted heterogeneity in the gene expression matrix by regressing out top surrogate variables while explicitly retaining temperature and *Wolbachia* effects (Leek and Storey, 2007). The numbers of surrogate variables were determined using the “num.sv” function in the “sva” package in R. The adjusted gene expression matrices were used for subsequent analyses for females and males separately.

### Quantitative genetic analysis of gene expression

Within each temperature and sex, we partitioned the observed phenotypic variance into genetic 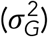 and micro-environmental 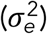 variances using mixed model implemented in the “lme4” R package, adding *Wolbachia* infection as a fixed effect. We then combined data from the two temperatures and partitioned the observed phenotypic variance into genetic and environmental components using different models as described below, which allowed us to make specific inferences. First, we tested variance homogeneity by likelihood ratio tests comparing models with either a single variance parameter or separate variance parameters for the environments. The models were fitted by the “nlme” R package and included temperature, *Wolbachia*, and the interaction between the two as fixed effects. Second, we also partitioned the observed phenotypic variance across environments into genetic 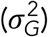, genotype by environment interaction 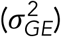, and micro-environmental variances 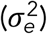, and the same fixed effects as above. We defined a GxE index as 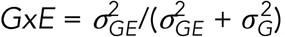, which measured the proportion of total genetic variance due to genotype by environment interaction. False discovery rate (FDR) was controlled using the Benjamini-Hochberg procedure when calling statistical significance. We compared genes significant for GxE with genes that showed evidence of adaptive divergence or GxE between tropical and temperate geographical locations, including genes that were differentially expressed between African and European populations (Hutter et al., 2008), differentially expressed between flies derived from Panama City and Maine at 21 and 29 °C (Zhao et al., 2015), and showed GxE among flies of temperate and tropical Australia raised at 18 and 30 °C (Levine et al., 2011).

### Gene set enrichment analysis (GSEA)

We performed quantitative GSEA at two instances with different gene specific scores. The first was a measure of canalization and decanalization and was computed as 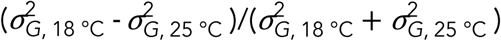. We considered only genes with at least one significant genetic variance at 18 or 25 °C. The second was a measure of GxE defined as 2GxE – 1 for genes with significant 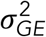 and/or 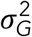. These scores were designed to range between −1 and 1 and replaced the correlation scores in GSEA, the general form of which takes a ranked list of scores as input. Subsequent steps were exactly followed as previously described in the original GSEA publication (Subramanian et al., 2005). We limited analyses to gene ontology terms with at least 20 genes among the entire set of genes with at least one GO annotation. For GO enrichment analysis for genes in Module 1 of the female WGCNA modules, the significance was tested using a hypergeometric test with genes in the WGCNA input as the background set. *P*-values were adjusted for multiple testing using the Benjamini-Hochberg procedure.

### eQTL mapping

We mapped eQTLs as previously described (Huang et al., 2015). Briefly, we obtained BLUP estimates of gene expression for each line at 18 °C, 25 °C separately, adjusting for *Wolbachia* infection, inversions, and major genotypic principal components (PCs). eQTLs were mapped using a *t* test implemented in PLINK. To estimate the empirical FDR, the phenotypic data were permuted 100 times, retaining the association between genes, and eQTLs were mapped following the same procedure. At each *P-*value threshold, the empirical FDR was estimated as the average number of significant eQTLs divided by the observed number. Significant eQTLs were further filtered by performing a forward selection procedure as previously described (Huang et al., 2015). In each iteration of the forward selection, all candidate eQTLs were tested conditional on eQTLs in the model, the one with the smallest *P-*value was added to the model until no eQTL could be added at *P* < 10^−5^. When considering matching of eQTLs between environments, an eQTL must be retained in the model selection in at least one environment. For example, an eQTL retained in the model selection at 18 °C may not be retained at 25 °C, but overlap is still assessed as long as it has been initially mapped at 25 °C before model selection.

### Co-expression network analysis

We identified modules in gene expression using the WGCNA R package (Langfelder and Horvath, 2008) and tested network preservation naively using the NetRep R package (Ritchie et al., 2016). To account for known cross-environment correlation of gene expression and quantitatively compare the preservation of correlation of expression among genes, we employed a simulation based approach. To test the null hypothesis that the plastic response of gene expression at 18 °C of different genes were independent from each other, we simulated for each gene its expression at 18 °C given the correlation between 18 and 25 °C. This was achieved by first scaling the expression at 25 °C such that *Z*_1_ = (*X* – *μ*_1_)/*σ*_1_, where *X* was the expression at 25 °C. Then the expression at 18 °C was simulated by the following formula 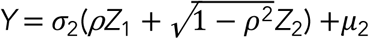, where *μ*_2_ and *σ*_2_ were the mean and variance of expression at 18 °C, *ρ* was the correlation coefficient between the expression at the two environments and *Z*_2_ was a random number drawn from the standard normal distribution. Thus *Y* was a simulated gene expression that preserved the mean and variance at 18 °C as well as the correlation and GxE between the two environments. We then computed the pair-wise correlation matrix at both temperatures and computed their difference. This procedure was repeated 1,000 times. A significant deviation from the null hypothesis was signified by decreased variance of the distribution of the correlation coefficient changes (Δ*r*).

We defined gene connectivity of a gene as the mean absolute correlation of the gene with all other genes. The strength of stabilizing selection was obtained from a previous study (Huang et al., 2016), in which mutational variance (*V*_*m*_) of gene expression was estimated in a set of mutational accumulation lines while the standing genetic variation (*V*_*g*_) was estimated from a subset of DGRP lines. The study also used whole bodies of flies of the same age but a different array platform and thus was an independent study from the present one. The strength of stabilizing selection was defined as the ratio of *V*_*m*_ over *V*_*g*_.

## Data availability

All data have been deposited into public repositories, including the RNA-Seq data (GSE67505 in GEO for 25 °C, PRJNA615927 in SRA for 18 °C), and the tiling microarray data in ArrayExpress (E-MTAB-3216 and E-MTAB-8953). All derivative data used to generate figures and tables are available in the GitHub repository https://github.com/qgg-lab/dgrp-plasticity-eqtl.

## Code availability

All codes used for analysis and figure and table generations are available in the GitHub repository https://github.com/qgg-lab/dgrp-plasticity-eqtl.

## Notes

### Competing Interest Statement

The authors have declared no competing interest.

